# Canagliflozin extends life span and leads to less weight gain in C57BL6 male mice

**DOI:** 10.1101/2022.11.20.517248

**Authors:** Evagelia E. Habeos, Fotini Filippopoulou, Menelaos Kanakis, George I. Habeos, George Lagoumintzis, Stavros Taraviras, Dionysios V Chartoumpekis

## Abstract

SGLT2 inhibitors are widely prescribed drugs for type 2 diabetes and heart failure. It seems that their beneficial health effects are multifaceted and not only limited to the amelioration of glycemic profile. It is suggested that SGLT2 inhibitors-induced glycosuria causes a metabolic shift that mimics the fasting response. It is also known that calorie restriction leads to enhanced longevity in mice. Thus, we hypothesized that long-term treatment of mice with SGLT2 inhibitors might extend their life span. To this end male C57BL6 mice at the age of 4 months were put on a normal chow diet or on a diet supplemented with 200 mg/kg canagliflozin. The canagliflozin-treated mice showed lower body weight gain over time and increased life span. The median survival of control mice was 107.5 weeks, while that of the canagliflozin-treated group was 112.5 weeks (p=0.011). No difference was seen in the presence or severity of cataracts. This study showed for the first time an enhanced median survival of canagliflozin-treated male mice with a homogeneous genetic background (C57BL6). Further analyses are in progress to elucidate the metabolic adaptations and mechanisms underlying this effect.

## 1. Introduction

Several studies have shown that caloric restriction can delay aging in mice and improve indices of health[1-3], with mechanisms suggesting growth hormone/IGF-1 signaling playing important roles [4]. Overall caloric restriction-induced longevity is associated with a reduction in the activity of growth pathways and a switch to metabolic patterns associated with fasting responses. These changes are coincident with reduced inflammation without a general impairment of immune function, which could contribute to protection against diseases ranging from cancer, to cardiovascular disease, to Alzheimer’s and autoimmune diseases [5]. Timing of onset of the diet, sex, and existing metabolic and genetic status can influence the efficacy of these diets in producing beneficial effects.

Several drugs that affect these pathways have been tested so far. In cohorts of genetically heterogenous mice, rapamycin [6], 17α-estradiol [7], nordihydroguaiaretic acid (NDGA) [8], protandim (14), glycine (17), acarbose [9] and aspirin [10] have been tested and showed extension of life span.

Several large cardiovascular or renal outcome trials of SGLT2 inhibitors have been carried out and showed benefit in patients with diabetes[11-13] or without diabetes[14, 15]. Although several mechanisms have been proposed [16, 17], it is generally believed that by shunting substantial amounts of carbohydrates into the urine, SGLT2 inhibitors-mediated glycosuria results in a progressive shift in fuel utilization toward fatty substrates and ketogenesis, partly through an increase in glucagon, and decrease in insulin levels [18-20]. The role of SGLT1 which has a broader distribution in the intestine and heart might add extra benefit[16]. Experiments in mice suggest that canagliflozin reprograms metabolism, modulates nutrient-sensitive pathways with activation of AMPK, and inhibition of mTOR, independent of insulin or glucagon sensitivity or signaling [21]. This metabolic shift is reminiscent of the fasting response that has been linked to prolonged longevity.

The aim of the study was to test the hypothesis that canagliflozin started early in life, might extend the life span in C57BL6 male mice. The selection of canagliflozin was based on the fact that besides SGLT2 inhibition, it inhibits to a lesser degree SGLT1, which is expressed in kidney, small bowel, and heart. It is also known that canagliflozin in C57BL6 mice promotes ketosis, and switch cell life programs by downregulating mTOR and upregulating AMP-activated protein kinase (AMPK) and fibroblast growth factor 21 (FGF21), resulting in energy conservation [21, 22]. During the current project’s execution another study came out that showed that canagliflozin extends the life span in genetically heterogenous male but not in female mice [23].

## 2. Materials and Methods

### 2.1 Mice

The study was conducted at the animal facility of the Medical School of the University of Patras. Mice were maintained in cages of 4 at 22°C ± 2°C on a 12-hour light/dark cycle, with *ad libitum* access to water and food. This animal protocol was approved by the Institutional Review Board Committee of the University of Patras, School of Medicine. Wild-type C57BL6J mice were initially obtained from Jackson Laboratory (Bar Harbor, ME) and were bred in our facility. Mice were crossed to generate male animals so as to enroll them in the study. As soon as they reached the age of 4 months, they were assigned either to control or canagliflozin group. The data presented herein were obtained from 83 male mice that consumed chow diet (control group) and 92 male mice that consumed chow mixed with canagliflozin at 200mg/kg (Cana group). Mice were followed until their death or when they developed severe signs of distress.

### 2.2 Mouse diet and treatment

Chow diet (product number ΒΠ-302) was purchased from Viozois, SA (Greece). Canagliflozin diet was custom made by the same company (Viozois, SA) using the same chow as basis and adding canagliflozin at 200 mg/kg concentration.The diet composition can be found in **Table 1** and the concentrations of additives can be seen in **Table 2**. Mice were inspected every week for their well-being up to 12 months of age, and then daily so as to document any diseased or dead animals. The carcasses of the deceased mice were preserved in buffered formalin for further evaluation. Body weights were recorded at the indicated time points. Retro-orbital sampling of blood was conducted for further analysis. Since it was not feasible to measure blood or tissue level of canagliflozin, urine was periodically tested for ketones using strips. In this way we confirmed the presence of urine ketones in Cana group.

**Table 1.** Diet composition.

| **Chemical composition** | **(%)** |
| --- | --- |
| Moisture | 12% |
| Crude protein | 18.5% |
| Crude fat | 5.5% |
| Crude fiber | 4.5% |
| Ash | 6% |
| NFE (nitrogen free extract) | 53.5% |

**Table 2.** Additives included in mouse chow.

| **Additives** | ***per kg of chow*** |
| --- | --- |
| Vitamin A | *15000 IU* |
| Vitamin D3 | *5000 IU* |
| Vitamin E (racemic mixture of all the stereoisomers of α-tocopherol) | *100 mg* |
| Vitamin K3 | *6 mg* |
| Vitamin B1 | *3 mg* |
| Vitamin B2 | *8 mg* |
| Vitamin B6 | *5 mg* |
| Vitamin B12 | *0.016 mg* |
| Calcium pantothenate | *30 mg* |
| Niacin | *85 mg* |
| Folic acid | *85 mg* |
| Biotin | *0.05 mg* |
| Vitamin C | *10 mg* |
| Iron (from iron sulfate) | *80 mg* |
| Manganese (from manganese oxide) | *120 mg* |
| Zinc (from zinc oxide) | *100 mg* |
| Copper (from copper sulfate) | *20 mg* |
| Iodine (from calcium iodide) | *1.5 mg* |
| Selenium (from sodium selenite) | 0.3 mg |
| butylated hydroxytoluene | 0.6 mg |
| Ethoxyquin | 0.6 mg |

### 2.3 Cataract evaluation

Cataract assessment was performed based on a slight modification of a previously described method [24]. Briefly, the presence of cataract was assessed with a direct ophthalmoscope by an expert ophthalmologist (MK) after applying a drop of atropine. Each eye was assigned a score from 0 (absence of cataract) to 5 (complete opacity of the lens with white marble-like appearance). In case of score discrepancies between the two eyes, the score of the highest scoring eye was assigned to the mouse.

### 2.4 Statistics

Student’s t-test was employed to assess the differences in body weights. Log-rank (Mantel-Cox) test was used to compare the survival curves. Mann-Whitney test was used to compare the severity of cataracts. GraphPad Prism 9 (GraphPad Software, La Jolla, CA, USA) was used for the statistical analysis and for the generation of graphs.

## 3. Results and Discussion

### 3.1 Canagliflozin-treated mice gained less weight over time compared to control ones

Mice were observed weekly for their well-being up to 12 months of age, and then daily so as to document any diseased or dead animal. At baseline (age of 4 months), control group (n=83) continued to consume regular chow while the treated group (n=92) switched to chow supplemented with 200 mg/kg canagliflozin. The two groups did not differ in their body weights at baseline (control body weight= 25.7 ± SEM 0.17 g versus canagliflozin group body weight= 25.6 ± SEM 0.2). After 10 months on the treatment, the canagliflozin group showed significantly less weight than the control (**Fig. 1**, control=33.1 ± SEM 0.26 g versus canagliflozin= 31.4 ± SEM 0.21g). Up to this point no deaths had been noticed in either group.

**Figure 1.**
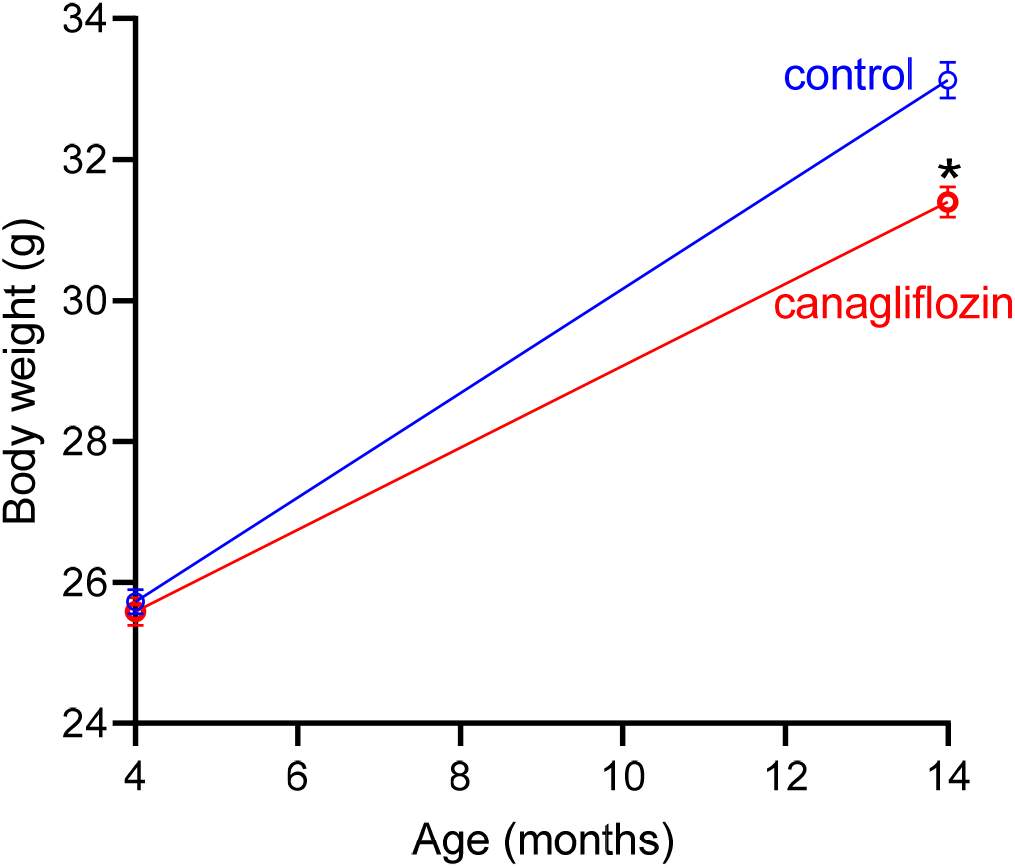
Body weights at the beginning of the treatment (age of 4 months) and after 10 months under treatment with canagliflozin or vehicle (age of 14 months). Data from 14 months onwards were omitted from this graph as mice started dying. Data show means ± SEM, *p<0.001 compared to vehicle control treatment at the same timepoint. N= 83 for control and n=92 for canagliflozin-treated mice. Student’s t-test was employed to assess differences at each timepoint.

### 3.2 Canagliflozin extends the lifespan of male C57BL6J mice

Mice were initially observed every week up to 12 months of age and later daily in order to record any diseased or dead animal. The survival curves of the control and the canagliflozin-treated mice diverged significantly, and the latter showed an improved survival (**Fig. 2**). Specifically, the median survival of control was 107.5 weeks while of the canagliflozin-treated group was 112.5 weeks (p=0.011).

**Figure 2.**
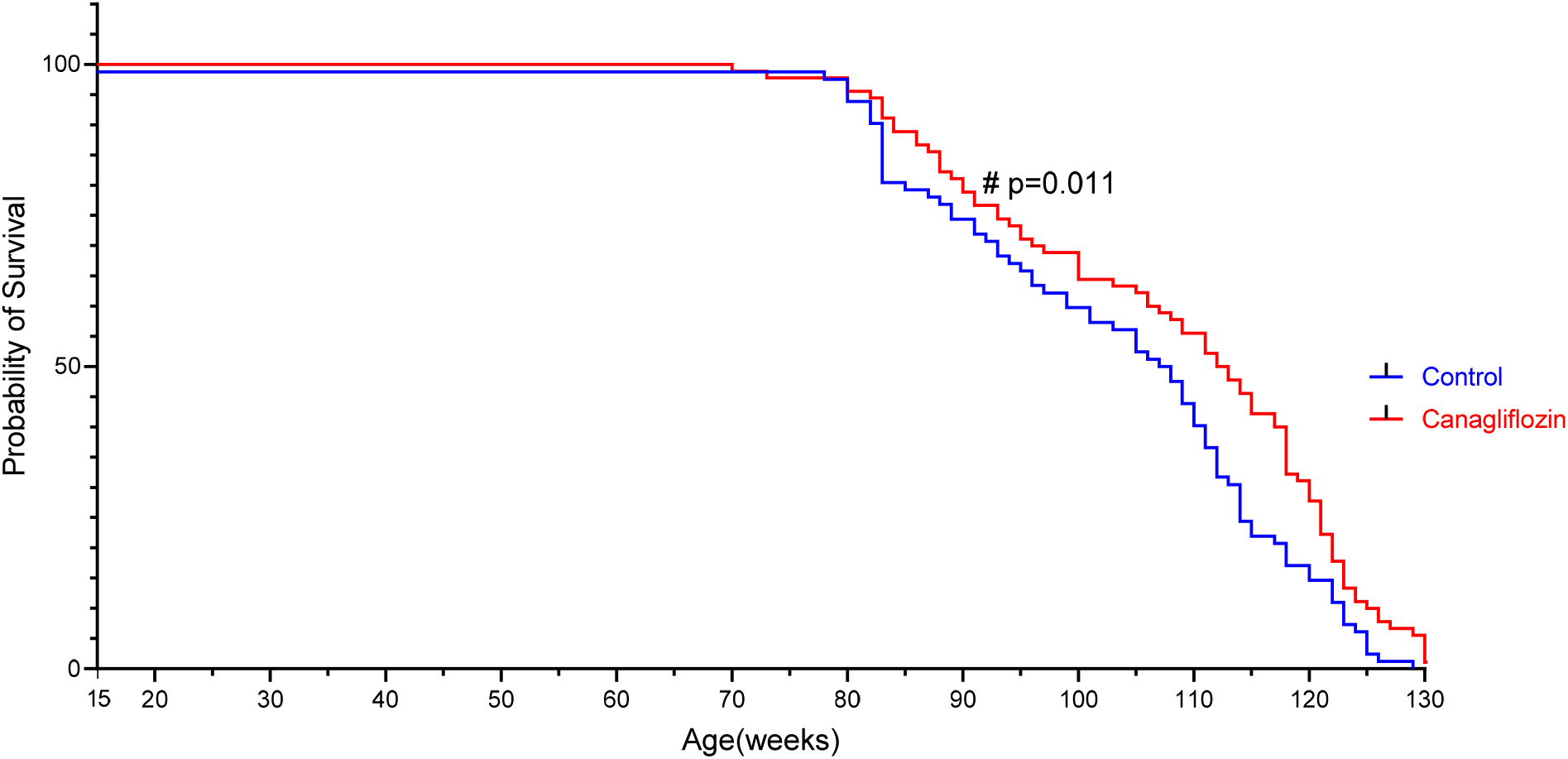
Survival curves of male mice fed a diet supplemented with canagliflozin or vehicle (control). The survival curves were compared using the Log-rank (Mantel-Cox) test. #p=0.011 signifies that the survival curves were different.

**Figure 3.**
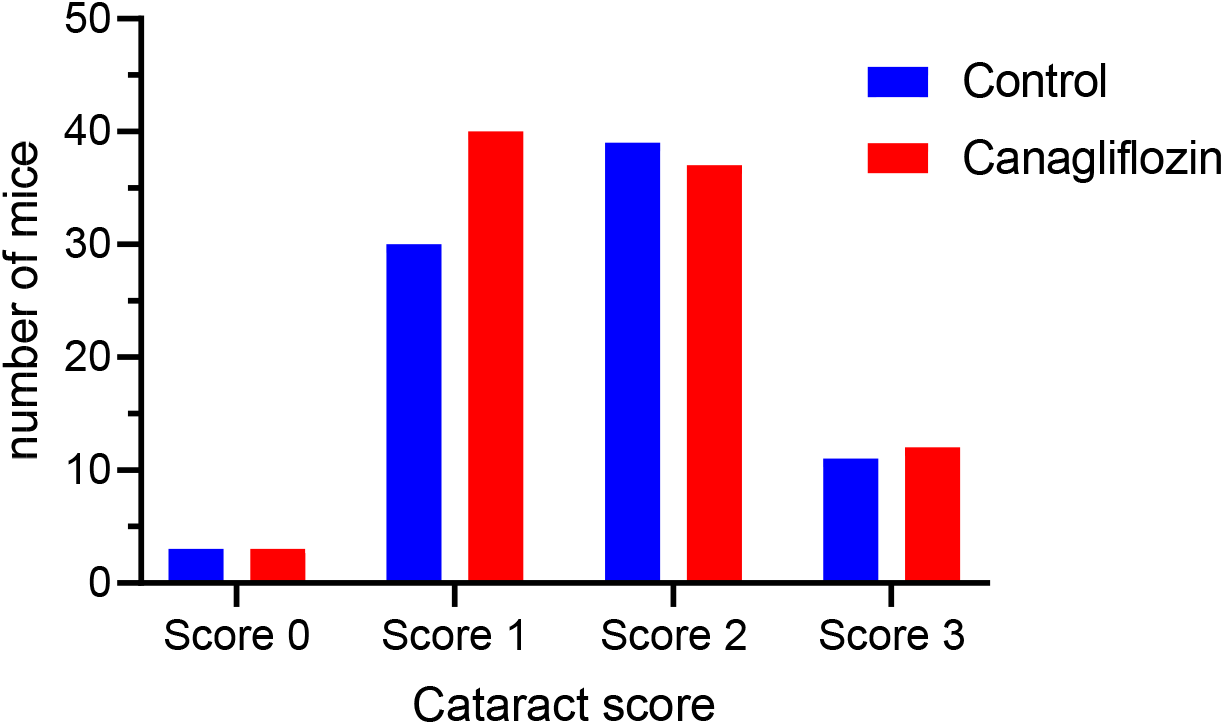
Cataract severity scoring in mice treated with vehicle (control) or canagliflozin at the age of 14 months. No statistically significant difference was found in the severity of cataract between the 2 groups. Mann-Whitney test.

This study was performed only in male mice; thus we cannot extrapolate these findings to female. This is a limitation of our study as sexual dimorphism has been noticed when assessing various interventions with reference to longevity. Specifically, a similar study in genetically heterogeneous mice demonstrated that canagliflozin increased median survival in male but not in female mice [23]. Our study differs from the aforementioned one since we used C57BL6J mice which have a genetically homogeneous background, are prone to metabolic diseases [25] and have been extensively used in various experimental settings.

#### Evaluation of presence and severity of cataract in control and canagliflozin-treated mice

C57BL6 mice are known to develop cataracts with age [26] Previous studies have shown that SGLT2 inhibitors can inhibit the progression of cataract in spontaneously diabetic Torii fatty rats [27, 28] and reduce the oxidative-stress induced markers in the lenses of fructose-induced diabetic rats [29]. We assessed whether canagliflozin treatment would have any impact on the development and/or severity of cataract. At the age of 14 months, no differences were recorded in either the presence or the severity of cataract between the two groups. Among the limitations of this assessment is its qualitative rather than quantitative nature, the use of an ophthalmoscope instead of a slit lamp and the lack of histology data [28].

## 4. Conclusion

This is a preliminary report of an ongoing study in male C57BL6 mice that were treated with canagliflozin since the age of 4 months. Canagliflozin treatment resulted in less gain of weight and increased median survival.

